# Amyloid precursor protein-b facilitates cell adhesion during early development in zebrafish

**DOI:** 10.1101/795187

**Authors:** Rakesh Kumar Banote, Jasmine Chebli, Tuğçe Munise Şatır, Gaurav K. Varshney, Rafael Camacho, Johan Ledin, Shawn M. Burgess, Alexandra Abramsson, Henrik Zetterberg

## Abstract

Understanding the biological function of amyloid beta (Aβ) precursor protein (APP) beyond its role in Alzheimer’s disease is emerging. Yet, its function during embryonic development is poorly understood. The zebrafish APP homologue, Appb, is strongly expressed during early development but thus far has only been studied via morpholino-mediated knockdown. Zebrafish enables analysis of cellular processes in an ontogenic context, which is limited in many other vertebrates. We characterized zebrafish carrying a homozygous mutation that introduces a premature stop in exon 2 of the *appb* gene. We report that *appb* mutants are significantly smaller until 2dpf and display perturbed enveloping layer (EVL) integrity and cell protrusions at the blastula stage.

Moreover, *appb* mutants surviving beyond 48 hpf exhibited no behavioral defects at 6 dpf and developed into healthy and fertile adults. The expression of the *app*-family members, *appa* and *aplp2*, was found to be altered in *appb* mutants. Taken together, we show that *appb* orchestrates the initial development by supporting the integrity of the EVL, likely by mediating cell adhesion properties. The loss of Appb might be compensated for by other *app* family members to be able to implement continued normal development.

## Introduction

The amyloid beta (Aβ) precursor protein (APP) is an integral membrane protein recognized for its role in Alzheimer’s disease (AD) pathogenesis. It is a single-pass glycosylated type I transmembrane protein that is processed by proteases into numerous extracellular and intracellular biologically active fragments (reviewed by (Andrew et al., 2016)). Cleavage of APP by β- and γ-secretases generates cytotoxic Aβ peptide fragments that aggregate into extracellular plaques in the brain in AD (Blennow et al., 2006; Hardy and Selkoe, 2002; Selkoe and Hardy, 2016). Moreover, recent studies have uncovered that APP might also contribute to AD-related neurodegeneration in Aβ-independent ways (Cheng et al., 2016; Pimplikar et al., 2010).

APP is ubiquitously expressed with especially strong expression in the central nervous system (Arai et al., 1991; Lorent et al., 1995; Selkoe et al., 1988). However, its basic biological role has been difficult to elucidate, likely due to redundancy between APP family members (von Koch et al., 1997). Nevertheless, studies suggest fundamental functions of APP in axonal transport, cell differentiation and proliferation, synaptic transmission, learning and memory, neuronal development, neurite outgrowth, as well as in cell adhesion (reviewed by (Deyts et al., 2016)). Mice with a deletion of *App* are viable and fertile but show reduced body weight, grip strength and locomotor activity (Müller et al., 1994; Zheng et al., 1995). Similarly, single *Aplp1* and *Aplp2* knockouts also display mild phenotypes. However, *App*^-/-^/*Aplp2*^-/-^ or *Aplp2*^-/-^/*Aplp1*^-/-^ double knockouts and triple knockouts (*App*^-/-^/*Aplp1*^-/-^/*Aplp2*^-/-^) show severe abnormalities and lethality (Heber et al., 2000).

Here, we have performed a detailed characterization of zebrafish with a homozygous mutation in *appb*, generated with CRISPR/Cas9 technology. As an animal model, zebrafish has several advantages over, *e.g.*, rodents; its external development, in combination with optical transparency, makes it possible to study cellular processes during early development in relation to genetic perturbation in a way that would be very challenging in mice. We analyzed morphological and behavioral changes, as well as potential compensatory mechanisms of other *app* family members, in *appb* mutants. During the blastula period, when zebrafish embryos transition from 128-cell stage to 50% epiboly, we observed enveloping layer (EVL) adhesion defects, of which the most severe were lethal. Our data suggest that this is caused by defects in cell adhesion. Similar to mice, mutant zebrafish were slightly smaller and did not show any behavioral alterations. However, loss of *appb* led to increased *appa* and *aplp2* expression, as determined by *in situ* hybridization and qPCR. Overall, these results suggest early and partially lethal abnormalities in the development of *appb* mutants, which in the majority of mutants may be prevented by increased expression of other *app* family members.

## Materials and methods

### Animal care and ethics statement

Fish were maintained in stand-alone racks on a 14h10h light:dark cycle at 28.5°C, at the facility of the Institute of Neuroscience and Physiology, University of Gothenburg. System water was kept at a pH of 7.2-7.5 and conductivity of 600μS. Larva were raised in Embryo medium (EM) (1.0mM MgSO_4_, 0.15mM KH_2_PO_4,_ 0.042mM Na_2_HPO_4,_ 1mM CaCl_2,_ 0.5mM KCl, 15mM NaCl, 0.7mM NaHCO_3_) at 28.5°C in dark except for those in behavior tests that were maintained at a 14h:10h light:dark cycle. This study was approved by the ethical committee in Gothenburg. All procedures for the experiments were performed in accordance with the animal welfare guidelines of the Swedish National Board for Laboratory Animals.

### Mutagenesis using the CRISPR/Cas9 system

The CRISPR/Cas9 system was used to generate zebrafish mutants, as previously described (Varshney et al., 2016). Briefly, gRNA synthesis was made with a cloning-free, oligo-based method using a target-specific DNA oligo (5’-GGATGACTCGGTGGGCTTGT-3’) and a ‘generic’ DNA oligo for the guide RNA. The two oligos were annealed and extended with DNA polymerase, and the resulting product serves as a template for *in vitro* transcription. Embryos were co-injected with 50pg gRNA and 150pg Cas9 mRNA, transcribed from the XbaI-linearized pT3TS-nls-zCas9-nls into the yolk at the one cell–stage. Injected embryos were screened for gRNA activity using a three-primer fluorescence PCR method. A 260bp region surrounding the target site amplified using M13F-tailed forward primer (5’-TGTAAAACGACGGCCAGTATTGAAGGATGCCCTCTTTG-3’) and a pig-tailed reverse primer (5’-GTGTCTTCCCATCGTGAAGCTGTCTAA-3’) in combination with a generic M13-FAM primer (FAM-5’-GTAAAACGACGGCCAGT-3’). The remaining embryos were raised to adulthood and outcrossed with wild-type fish. Seven to eight embryos from each outcrossed pair were screened for mutations in the F1 generation. Two different alleles were selected: *appb*^*26_2*^ (−5bp) and *appb*^*26_4*^ (−8bp). These carry frameshift mutations and were raised and subsequently outcrossed towards the wild-type AB fish line until generation F4. Outcrossed adults were genotyped using an allele-specific KASP assay (LGC, Middlesex, UK). Offspring from heterozygous F4 inbreeds were used to generate homozygous wild-type and mutant lines. Embryos/larvae from these adult zebrafish were used to eliminate the influence of other potential genetic differences.

### Body length measurement

Microscopic images were acquired using a Nikon stereomicroscope (Mellville, NY, USA). For the one-cell stage, the size of the embryo was measured across the yolk. The total body length of 24 hpf, 48 hpf and 3 dpf embryos was measured from the frontal part of the head to the end of the tail fin, along the anterior-posterior axis. Scales at the same magnification as the images were employed in the measurement.

### In situ hybridization

Antisense digoxigenin-labeled RNA probes were generated from linearized DNA templates against *appb* (Abramsson et al., 2013; Banote et al., 2016) and *aplp2* (MDR1734-202739496, Open Biosystems). *Appa* riboprobes were generated by amplifying fragments from cDNA clones, inserting them into the pCR2.1-TOPO vector (K450001, Invitrogen), linearizing the vector and transcribing with RNA polymerase using the following oligonucleotides: *appa*, Fwd 5’-AGGCGCATCGCGTTCTTCACAGAG-3’ and Rev 5’-CCTAACCCTCCCCGAACCCTCCC-3’. The *aplp1* riboprobe was transcribed from PCR-amplified fragment using primers with T7 or T3 linkers (Fwd *5’-* TAATACGACTCACTATAGGGTCGCGGTGTGGAATATGTCTGCT-3’ and Rev 5’-GCAATTAACCCTCACTAAAGGGTCTGCCTCTGCCCACTCCTTCA-3’). Whole mount RNA *in situ* hybridization (WISH) was performed as described earlier (Andermann et al., 2002) with modifications (Banote et al., 2016).

### Immunofluorescence and confocal microscopy

Embryos at sphere (4 hpf) and germ ring (5.7 hpf) stages were fixed overnight in 4% paraformaldehyde at 4°C and then washed sequentially in phosphate-buffered saline (PBS), 0.5% Triton-X in PBS (PBT) and then dechorionated. Embryos were then incubated in block solutions (5% normal goat serum, 1% bovine serum albumin [BSA], 1% DMSO and 0.5% TritonX-100 in PBS) for 2 hrs and incubated overnight at 4°C in block solution containing 1:50 Phalloidin Alexa 568 (A12380, Molecular Probes) to stain actin, or antibody towards E-cadherin (610181, BD Bioscience), β-catenin (C2206, Sigma-Aldrich) or ZO-1 (33-9100, ThermoFisher Scientific). The next day, embryos were washed gently in PBT and incubated overnight (ON) with secondary antibody with the desired fluorescent conjugate or were mounted in 1% agarose in glass bottom dishes (D29-10-1.5-N, Cellvis) in PBT medium and imaged using Zeiss LSM710 confocal microscope (Carl-Zeiss, Jena, Germany) using 40x water immersion objective (Plan-Apochromat 40x/1.0) with 488 and 568nm laser.

For proliferation assay, wild-type and mutant embryos at 4 hpf were fixed in 4% PFA for 1.5 hrs at room temperature, washed with PBS and incubated with ice cold 100% acetone at −20°C for 20-30 min. Samples were transferred to a 24-well plate and washed five times with PBT on gentle agitation. Nonspecific binding was blocked with block solution for at least 60 min and then incubated over night at 4°C with primary antibody against phospho-Histone3 (pHH3, 06-570, Millipore) diluted 1:500 in block solution. Samples were washed carefully 5×10 min in PBST at RT and incubated with goat anti-rabbit-Alexa488 (A-11008, Molecular probes) over night at 4°C or 2 hours at room temperature. Samples were carefully washed five times for 10 min in PBST. During final wash, DAPI (D1306, Molecular Probes), 1:1000 and Alexa Fluor 568 Phalloidin (A12380, Molecular Probes) 1:300 were added to the PBST wash solution. Blastulas were mounted in 1% low melting agarose on glass bottom 35 mm Petri dish (81158, Ibidi). Stacks were acquired with an inverted Nikon A1 confocal system (Nikon Instruments, Mellville, NY, USA) using a 20x objective. Quantification of proliferating EVL cells was performed manually by counting phalloidin- and pHH3-positive cells located at the outer cell layer of the blastula using ImageJ software (National Institute of Health, USA).

### Quantification of cell shape /morphology

Confocal images of phalloidin-stained embryos were analyzed by ImageJ 1.52a software (National Institute of Health, USA). Images were converted to binary and the threshold set manually to visualize cell borders. Noise was removed with Despecle plugin. Cell borders were outlined in binary images with the Watershed plugin. False borders were manually adjusted. Cell number, cell area and cell circularity were analyzed with the plugin Analyze particles. Junctions were manually counted. Image analysis were performed in a blinded fashion on decoded images.

### Antibody production

A polyclonal antibody was produced by immunizing rabbits with a peptide corresponding to the N-terminal half of the zebrafish (Aβ) fragment (Ac-EERHNAGYDVRDKRC-CONH_2_) (Agrisera, Umeå, Sweden), since this sequence differs significantly between Appa and Appb. Antibodies from the second bleed were protein G-purified.

### Western blotting

Adult brains or embryos at 24 hrs were deyolked by pipetting in embryo medium. Brains and sixty embryos were homogenized with a 23G syringe and lysed in lysis buffer (10 mM Tris-HCl pH 8.0, 2% sodium deoxycholate, 2% SDS, 1 mM EDTA, 0.5 M NaCl, 15% glycerol) supplemented with protease inhibitor cocktail (04693132001, Roche). Samples were sonicated for 10 min and incubated on ice for 20 min. Supernatants were collected and protein concentration was measured using the BCA Protein Assay Kit (23225, ThermoFisher Scientific). Proteins were separated on a NuPAGE™ Novex™ Bis-Tris pre-cast gel (Invitrogen) and transferred onto 0.2 μm nitrocellulose membrane (GE Healthcare). The membrane was blocked with 5% milk and immunoblotted with anti-APP (Y188) antibody (ab32136, Abcam) at a dilution of 1:2000 (100% homology to zebrafish Appb and 93% to Appa) or the G-protein purified in-house generated polyclonal rabbit anti-Appb antibody generated against the N-terminal half of the Aβ-peptide. Immunoreactivity was visualized by anti-rabbit HRP-linked secondary antibody (7074S, Cell Signaling) at 1:5000 dilution. The signal was developed using SuperSignal West Dura Extended Duration Substrate kit (Thermo-Fisher) and imaged using ChemiDoc Imaging Systems (Bio-Rad). Thereafter, blots were re-probed with 1:20000 GAPDH antibody (2D4A7) [HRP] (Novus Biologicals) or mouse anti-αTubulin (T6199, Sigma-Aldrich) as a loading control. Western blot images were analyzed, produced and quantified using Image Lab™ Software (Bio-Rad). The intensity of the App band was related to glyceraldehyde 3-phosphate dehydrogenase (Gapdh) band intensity in a ratio.

### Quantitative PCR

A total of 40 embryos were split into 10 embryos per sample and total RNA was extracted using TRI Reagent^®^ (T9424, Sigma-Aldrich). Then, cDNA was synthesized using High Capacity cDNA kit (Applied Biosystems) with RNase inhibitor and converted in a single-cycle reaction on a 2720 Thermal Cycler (Applied Biosystems). Quantitative PCR was performed with inventoried TaqMan Gene Expression Assays with FAM reporter dye in TaqMan Universal PCR Master Mix with UNG. The assay was carried out on Micro-Amp 96-well optical microtiter plates on a 7900HT Fast QPCR System (Applied Biosystems). qPCR results were analyzed with the SDS 2.3 software (Applied Biosystems). Briefly, cDNA from each sample was normalized with average C_T_:s of eef1a1l1 and actb1, then the relative quantity was determined using the ΔΔC_T_ method (Livak and Schmittgen, 2001) with the sample of wild-type sibling embryos (24 hpf) as the calibrator. TaqMan^®^ Gene Expression Assays (Applied Biosystems) were used for the following genes; Amyloid Beta (A4) Precursor Protein A (*appa*; Dr 031 443 64_m1), Amyloid Beta (A4) Precursor Protein B (*appb*; Dr 030 803 08_m1), Amyloid Beta Precursor Like Protein 1 (*aplp1*; AJCSWD2), Amyloid Beta Precursor Like Protein 2 (*aplp2*; Dr03437773_m1), Eukaryotic Translation Elongation Factor 1 Alpha 1, Like 1 (*eef1a1l1*; Dr 034 327 48_m1) and Actin, Beta 1 (*actb1*; Dr 034 326 10_m1).

### Ionomycin treatment

Ionomycin (I0634, Sigma-Aldrich) was dissolved in EM prior to exposure. Wild-type and *appb* mutant embryos were treated with 5μM ionomycin at the 32-cell stage (1.75hpf) and compared at the blastula stage (2.5hpf). DMSO (0.1%) in EM was used as a control.

### Cell dissociation and aggregation assay

Dissociation and aggregation of blastoderm cells were performed as described in (Montero et al., 2003) with some modifications. Embryos were injected at the one-to four-cell stage with 10μg/μl of 10’000 Mwt dextran-Alexa Flour 488 (D22910, Molecular Probes) or tetramethylrhodamine-dextrane (D1824, Molecular Probes) diluted in 0.2M KCl and filtered through a 0.22 μm filter. At 4 hours post fertilization, embryos were manually dechorionated on an agarose plate and transferred to a deyolking buffer (55 mM NaCl, 1.8 mM KCl, 1.25 mM NaHCO_3_). Embryos were dissociated by pipetting up and down using a glass Pasteur pipette. Blastoderm cells were harvested by centrifugation at 400xg for 3 min at RT. Cells were washed twice with sterile DPBS (14190086, Invitrogen), resuspended in L15 medium (11415049, ThermoFisher), containing 2mM L-glutamine (G7513 Sigma-Aldrich, Munich, Germany), 100U/ml Penicillin, 100μg/mL Streptomycin (P4333, Sigma-Aldrich), 10% FBS and 1.25mM CaCl_2_. The cell suspension was transferred through a 40 μm Cell strainer (734-0002, Corning) and transferred to a glass bottom 35mm Petri dish (81218, LRI Instrument AB,) to a final concentration of 50 embryos/ 35mm dish. Petri dishes were coated with 10μg/ml fibronectin (11051407001, Sigma-Aldrich) at 37°C for at least 2h and washed with DPBS to remove excessive fibronectin. Cell cultures were incubated at 31°C in an incubator or heating chamber on the confocal microscope while cluster formation was analyzed. Confocal planes were taken at 1, 6 and 16 hours post dissociation (hpd) with a 10x objective (Plan Apochromat 10x/0.45NA) on a Nikon A1 Confocal Microscope System (Nikon Instruments, Mellville, NY, USA). Three separate experiments (*n*=3) performed on larvae from different parents were used for image analysis.

#### Analysis of aggregation and segregation

Collected images were treated as a matrix of dimensions x,y,z, where x is the number of row pixels, y is the number of column pixels and z is 2 - the number of color channels (red and green). To automatically segment the aggregates in the field of view, maximum intensity z-projection images were processed in FIJI software as such: contrasts were enhanced via the Normalize Local Contrast method. Smoothing was done via a Gaussian Blur of sigma equals 1. Image thresholding was done via Li’s Minimum Cross Entropy method. Each aggregate was then identified as a proper region of interest (ROI) and objects smaller than 1000 pixels were removed from the segmented image (software tools for segmentation and quantification are available at : https://github.com/CamachoDejay/). To characterize the segregation of the aggregates, we implemented a method previously described (Schötz et al., 2008). In brief, P (the electrical dipole moment) and S (the moment of inertia and ratio of scattering amplitudes) parameters were calculated. For this, each image color channel, red and green, was loaded as a separate matrix, where the matrix value indicated intensity and the row and column index represented the pixel position (intensity matrix l[r,c], where ‘r’ and ‘c’ are row and column indices). A full description of S and P calculations can be found in supplementary materials (Supplementary data 1). Data from three technical repeats (*N*=3) of clusters from a total of wt^(G)^/wt^(R)^ (*n*=742) or wt^(G)^/mut^(R)^ (*n*=1161) were plotted in GraphPad.

### Behavioral analysis

Adult heterozygous *appb* carriers (F3) were inbred to facilitate the study of locomotion of wild-type (*n*=30) and mutant (*n*=23) siblings. Larvae were incubated at 28.5°C on a 14:10h light to dark cycle. At 6 dpf, larvae of mixed genotypes were placed in 1mL EM in a 48-well plate and acclimatized overnight in the behavior room at 26°C and subsequently 20 min in the ZebraBox before initiation of tracking. Locomotor activity was recorded using the ZebraBox tracking system (ViewPoint, Lyon, France), equipped with infrared digital camera. Recording was carried out for 60 min in continuous light. Larvae were genotyped with a KASP assay (LGC, UK) specific for the mutation. The experiment was repeated twice. Swimming behavior was quantified using ZebraLab™ software (ViewPoint, Lyon, France), values were calculated in Microsoft Excel and graphs were plotted in GraphPad Prism.

### Statistical analysis

Statistical analysis was performed using GraphPad Prism® 7 software. Data were presented as means and standard errors of the mean (±SEM). For body length analysis, western blot analysis, locomotor activity over 60 min, qPCR data, cell shape, proliferation and analysis of scattering and dipole moment were performed with unpaired Student’s *t*-tests. Statistical significance was set at *p*<0.05.

## Results

### appb mutants show morphological defects at early stages

We established two *appb* mutant lines with the CRISPR/Cas9 method. The gRNA construct targeting the 5’-end of exon 2 (depicted with an arrow, Fig. 1A) generated a deletion of five nucleotides (dashed line) and insertion of two nucleotides (lower case letters) referred to as *appb*^*26_2*^ and a deletion of eight nucleotides as in *appb*^*26_4*^ (Fig. 1B). Sanger sequencing was used to confirm the mutations and showed disrupted reading frame and a premature stop codon at the 3’-end of exon 2 (amino acid 55, indicated by a black dot), as represented by the chromatograms (Fig. 1C). Both alleles showed the same phenotype and we therefore used allele *appb*^*26_2*^, hereafter referred to as the *appb* mutant.

**Fig 1.**
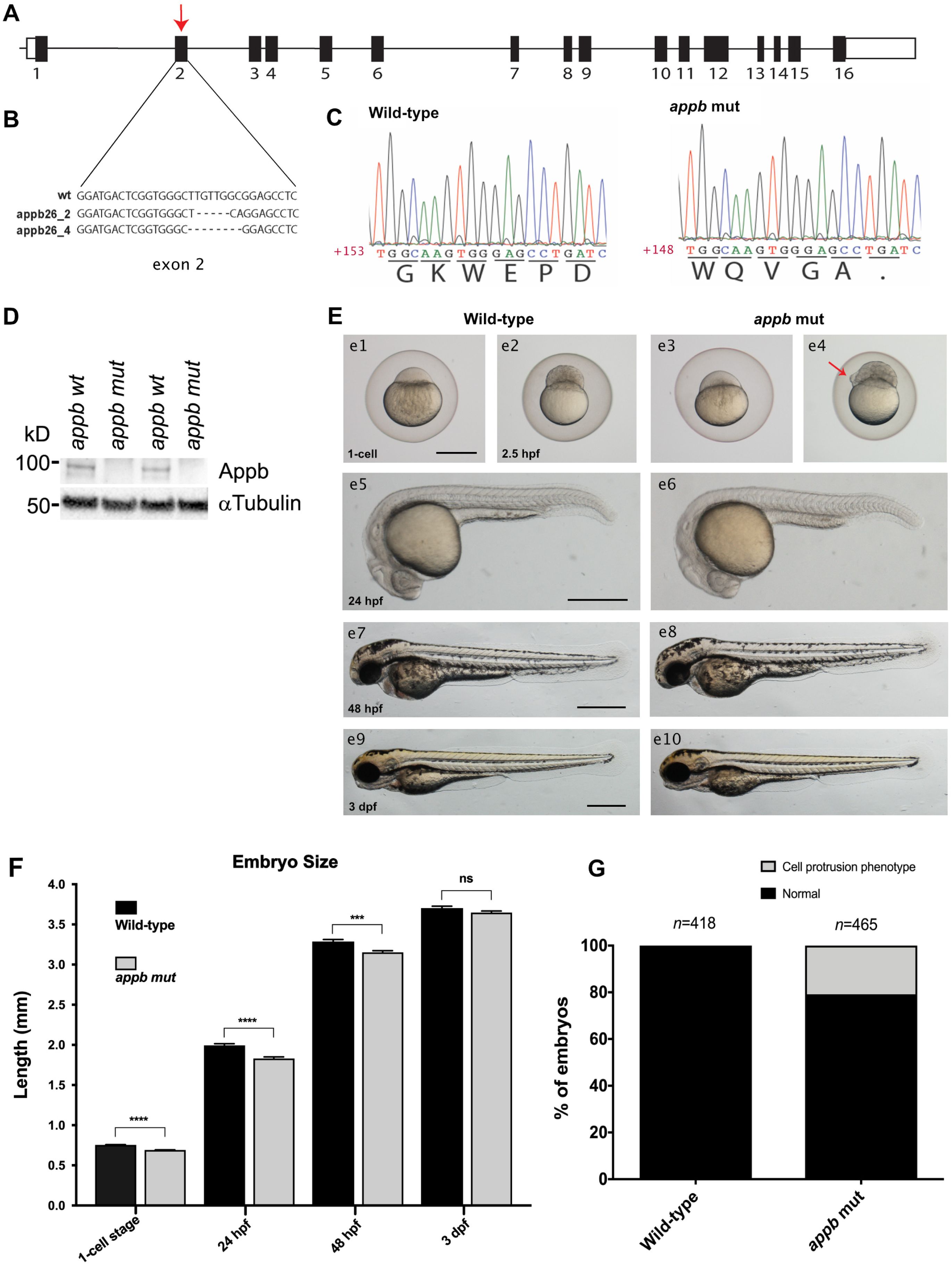
Characterization of the *appb* mutation. (A) Schematic representation of the coding and non-coding regions of the *appb* gene, indicating the mutation point (pointed arrow in red). (B) Schematic illustration of nucleotide sequence and deletion (dashed line). (C) Sanger sequencing chromatogram of the premature stop in exon 2. (D) Western blot analysis of Appb expression, *n*=2. (E) Morphology of *appb* mutants and wild-type zebrafish embryos from one-cell stage to 3 dpf larvae (e1, e3, e5, e7, e9 indicates wild-type and e2, e4, e6, e8, e10 are *appb* mutants). (F) Embryo size measurement at 1-cell stage, 24 hpf, 48 hpf and 3 dpf. (G) Quantification of disturbed cell adhesion phenotype. Scale bar= 500μm.

Next, we wanted to examine the effect of the *appb* mutation on protein expression by western blot. We produced a polyclonal antibody towards the N-terminal part of the Aβ peptide, since we could not find any good commercially available antibodies that were specific to the zebrafish Appb protein. The N-terminal part of the Aβ region shows low homology between Appb and the other members of the App family and was therefore considered as a good candidate peptide to use for immunization. While protein with the expected size was found in wild-type brains, no bands were detected in *appb* mutant brains (Fig. 1D). α-Tubulin, used as a loading control, verified equal protein loading in the different lanes. These results show that the mutation result in a loss of Appb protein expression.

Next, we performed gross morphological analysis of *appb* mutants from the 1-cell stage to 3 dpf larvae. The *appb* mutants, generated from homozygous parents, were significantly smaller at the 1-cell stage compared with wild-type embryos and had shorter body length until 48 hpf (Fig. 1E: e1, e3, e5, e6, e 7, e8; and Fig. 1F). In addition, we observed cells protruding through or perching on the EVL in ∼20% of mutant blastulas (Fig. S2, 1E: e2, e4, arrow and G), out of which some resulted in abnormal blastula formation (Fig. S1). This phenotype was detected around 2.25 hpf when the EVL just has formed. While ∼4% of these embryos later died, the remaining embryos developed further and survived, although with a slight delay in epiboly (Fig. 2). Intriguingly, no other gross morphological changes were observed at later stages compared with wild-type fish (Fig. 1 E: e9-e10 and Fig. 1F). Even though most embryos somehow managed to overcome this perturbation to develop into healthy and fertile adults, we hypothesized that the early defects might indicate failing EVL integrity in mutants.

**Fig 2.**
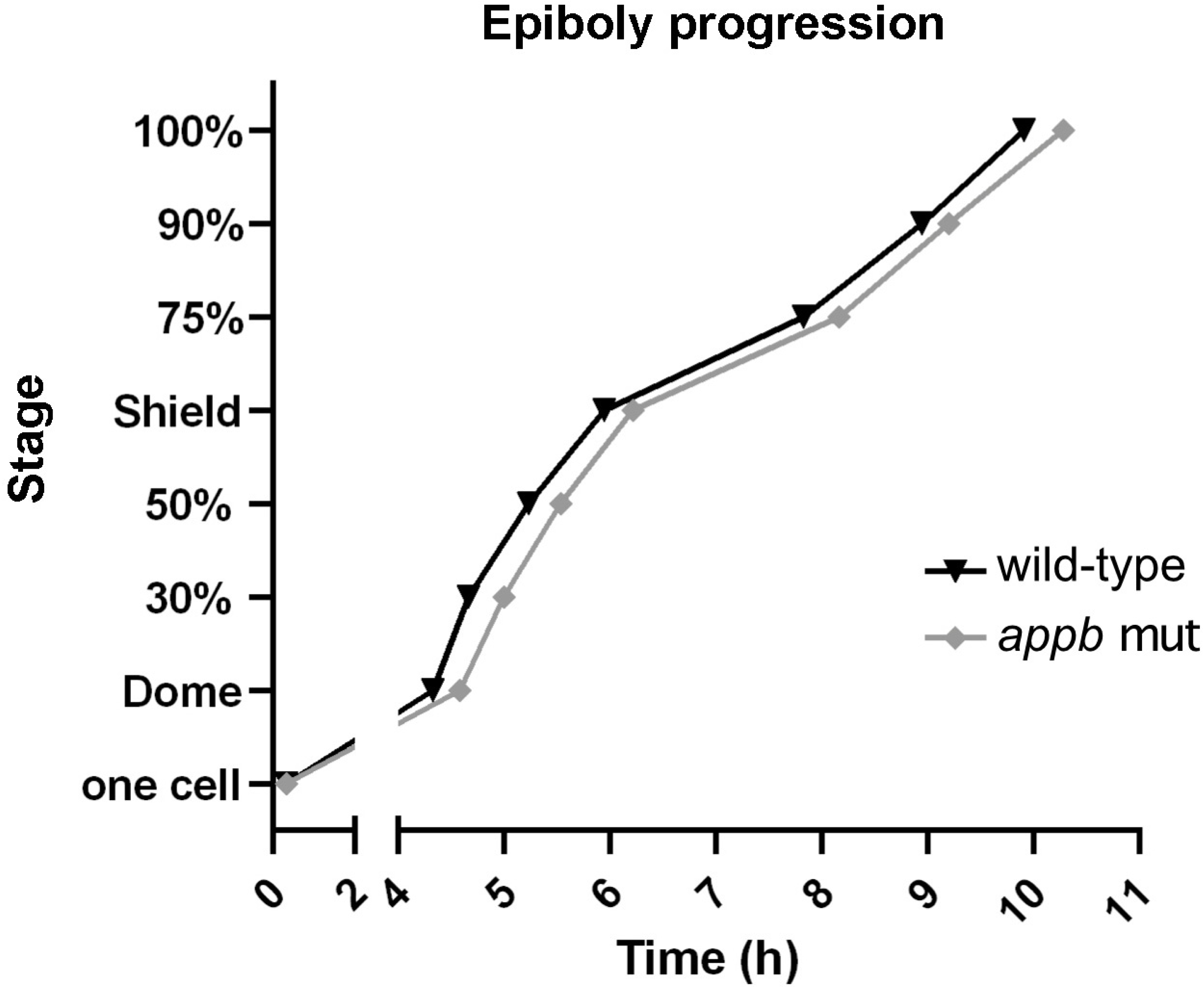
Delayed epiboly progression in *appb* mutants. The time (h) for wild-type (*n*=30) and *appb* mutant (*n*=30) zebrafish to reach dome stage and epiboly progression, measured as percentage, was analyzed. Staging was initiated when embryos reached one-cell stage.

### Defects in EVL integrity

Similar phenotypes, including protrusions and delayed epiboly, have been previously described in other mutants with EVL defects (Eno et al., 2016; Slanchev et al., 2009; Song et al., 2013a; Vannier et al., 2013). We therefore continued with analyzing the cellular arrangement of the EVL and performed phalloidin-staining of actin organization at the sphere (4 hpf) and germ ring stages (5.7 hpf), as these stages are easy to distinguish and thus allow for accurate staging (Fig. 3A,B). Interestingly, EVL cells in *appb* mutants had a larger cell surface area and were less circular compared with EVL cells of wild-type blastulas at 4 hpf (Fig. 3D, E). At the sphere stage, the cell cycle of the EVL slows down significantly (Kimmel et al., 1995) and the EVL is suggested to spread mainly by cell flattening (Campinho et al., 2013). However, if the EVL consists of fewer cells, the remaining cells would need to stretch more to cover the same area. We therefore analyzed whether the increase in EVL cell surface in mutants was due to decreased proliferation and thus changes in EVL cell number. However, we could not find any difference in the total number of EVL cells (Fig 3C) nor in the number of EVL cells going through mitosis by staining wild-type (Fig. 4A,C) and mutant (Fig. 4B,C) blastulas for phospho-histone 3 (pHH3), DAPI (nuclei) and phalloidin (F-actin) at 4 hpf.

**Fig 3.**
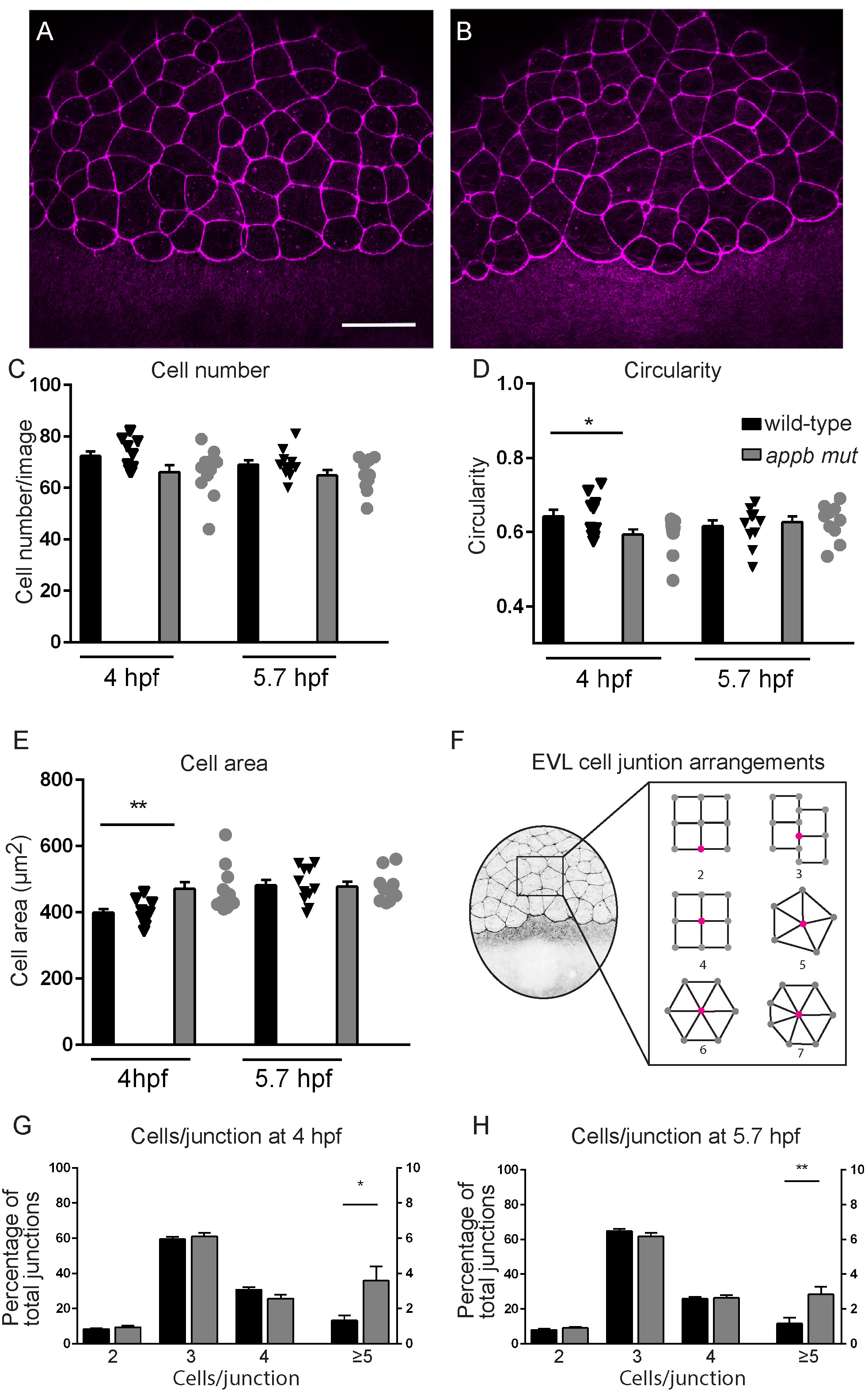
Changes in cell shape morphology in *appb* mutants. (A) Wild-type and (B) mutant embryos stained with phalloidin at 4 hpf. (C,D,E) Quantification of cell number and morphology. (F) Schematic of embryo at blastula stage and junction points connecting to number of cells. (G,H) Quantification of cells per junction points at sphere and germ ring stage. (Sphere: *n*=12 wt, 12 *appb* mut and germ ring: *n*=11 wt, 10 *appb* mut).

**Fig 4.**
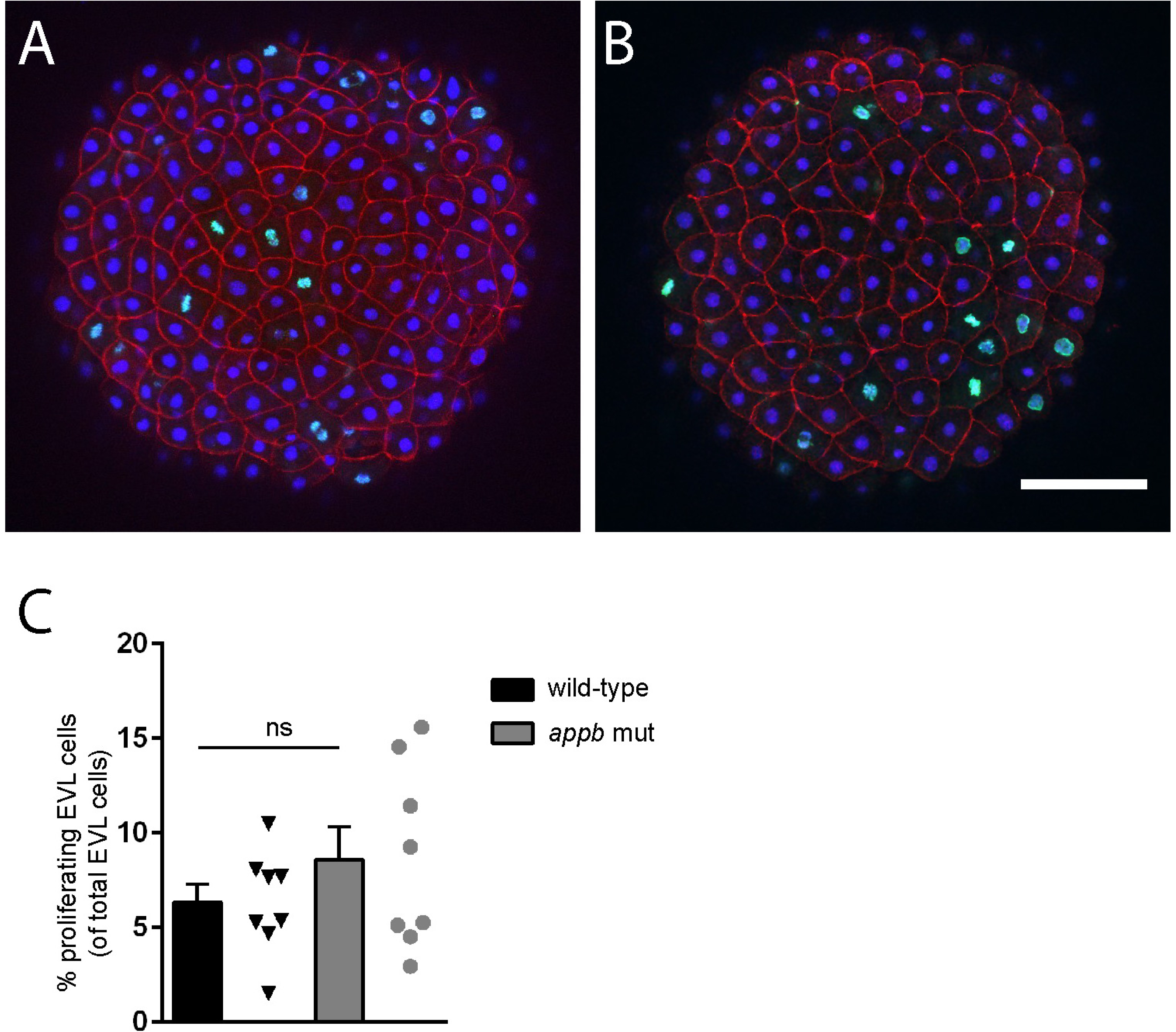
Proliferation of EVL in wild-type and *appb* mutants at 4 hpf. Confocal images of the EVL in wild-type (A) and *appb* mutant (B) embryos immunostained for the mitotic marker phospho-Histone H3 (pHH3; green) and actin binding phalloidin (red) at 4 hpf. All nuclei were labeled with DAPI (blue). Number of pHH3-positive nuclei located at the EVL surface (C). Student’s *t*-test was performed on wild-type (*n*=8) and *appb* mutant (*n*=8). Scale bar, 100μm.

The process of epiboly, starting shortly after sphere stage, is an orchestrated event depending on tissue expansion and radial intercalation of the blastoderm (Morita et al., 2017). These processes are proceeding under surface pressure, coordinated by the cells of the EVL through cell rearrangements and cell divisions(Morita et al., 2017). Thus, to understand the consequence of changed EVL morphology on cell arrangement, we quantified the number of cells per junction point, where a junction point was defined as the intersection between several cell membranes. (Fig. 3F, diagrammatic representation). In the wild-type blastula, at 4 and 5.7 hpf, we observed, most frequently, 3 or 4 cells sharing one junction point (Fig. 3G, H). A smaller percentage of all junctions were shared by only two cells. These were mainly present at the EVL margin close to the yolk syncytial layer. These EVL junctional formations were similar between *appb* mutants and wild-types (Fig. 3G; H). In addition, we found rosette-like junction arrangements with 5 cells in wild-types. However, mutants had significantly more rosette-like formations with up to 7 cells connecting to one junction point (Fig. 3G, H). Together, these results show that mutants have abnormal EVL morphology and arrangement. Changes in shape might be affected by defects in cell adhesion (Slanchev et al., 2009), and since both APP and APLP2 have been shown to be adhesion molecules at synapses (Soba et al., 2005), we hypothesized that cell-shedding together with changes in cell shape might reflect failing cell-cell adhesion.

### appb mutants are sensitive to ionomycin treatment

To investigate if the observed perturbations were a consequence of defective cell-cell interactions, we tested if mutant embryos were more sensitive to additional deterioration of cell adhesion. Ionomycin is a Ca^2+^ influx stimulator that promotes shedding of the adherens junction proteins cadherins (Esterberg et al., 2013; Marambaud et al., 2002), which leads to cell-cell detachment. We hypothesized that although not all mutant embryos displayed cell protrusions and shedding, they might still be more sensitive to ionomycin exposure. We treated wild-type and *appb* mutants with 5μM ionomycin at the 32-cell stage (1.75 hpf), observed them at the blastula stage (2.5 hpf) and quantified the phenotypes based on severity (Fig. 5). Wild-type embryos treated with control DMSO showed no phenotype, while ∼17% of mutant embryos showed cell adhesion phenotypes similar to incubation in EM (Fig. 5A, B, E). Interestingly, we found that around 77% of mutant embryos were severely affected/dead by the ionomycin treatment (Fig. 5D, E) compared with 10% of ionomycin-treated wild-type embryos (Fig. 5C, E). These data indicate that mutant embryos are more sensitive to additional weakening of cell-cell adhesion compared with wild-type embryos, supporting the hypothesis that Appb affects cell-cell interactions either directly or through other adhesion molecules.

**Fig 5.**
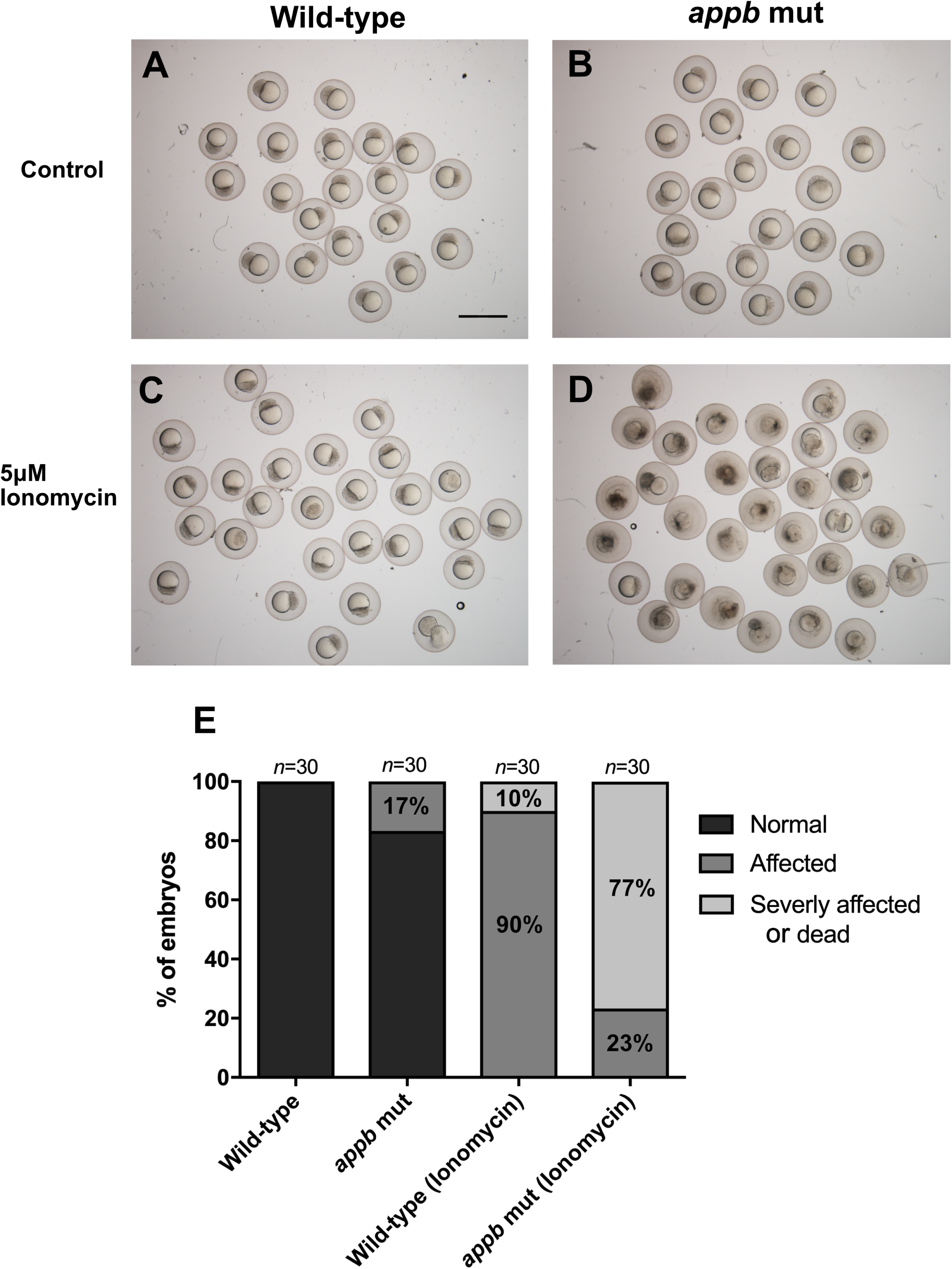
Increased sensitivity to ionomycin treatment in *appb* mutants. (A,B) Wild-type and *appb* mutant embryos treated with DMSO control. (C,D) 5uM ionomycin treatment of wild-type and *appb* mutants embryos. (E) Quantification of ionomycin sensitivity. Scale bar, 1500μm.

### Appb controls cell adhesion

Differences in cell-cell adhesion can be evaluated by an aggregation test where dissociated cells are allowed to re-aggregate *in vitro*. In such experimental setup, cells with different adhesiveness will segregate from each other, while cells with equal adhesiveness will mix randomly (Davis et al., 1997; Montero et al., 2003; Schötz et al., 2008; Steinberg, 1970; von der Hardt et al., 2007). To address if the adhesive property of *appb* mutant cells are different from wild-type cells, we therefore carried out a cell dissociation and re-aggregation test. Wild-type and mutant embryos were labeled by injections of red (R) or green (G) fluorescent dextran at the one cell stage, dissociated and plated at a 1:1 ratio in combinations of wt^(G)^/wt^(R)^ and wt^(G)^/mut^(R)^ (Fig. 6A,B). Although, aggregates started to form at 6 hours post dissociation (hpd) (Fig. 6C,D), the segregation process was allowed to proceed until 16 hpd (Fig. 6E,F). At this stage, both wild-type and *appb* mutant cells adhered to form well-defined clusters. Clusters containing wt^(G)^/wt^(R)^ cells (*n*=742) or wt^(G)^/mut^(R)^ cells (*n*=1161) were intermixed and did not show the clear segregation patterns as previously reported between cells with large differences in adhesion (Lachnit et al., 2008; Schötz et al., 2008; Song et al., 2013a; Vannier et al., 2013). On the other hand, the mild phenotype of *appb* mutants predicted a less dramatic change in the segregation process. Thus, to analyze more subtle changes in cell distribution, we took advantage of the image analysis method developed by Schötz and colleagues (Schötz et al., 2008) that use two parameters to quantitatively define scattering (S) and dipole moment (P) of cells within clusters. S is defined as S=S_red_/S_green_ where an S close to 1 represent intermixed cells and close to 0 or 2 when one cell group is segregating from the other. On the other hand, P describes separation of two populations and is close to 1 if cells in a cluster are completely separated into a red and green compartment but close to 0 when cells are fully mixed or when one cell type surround the other.

**Fig 6.**
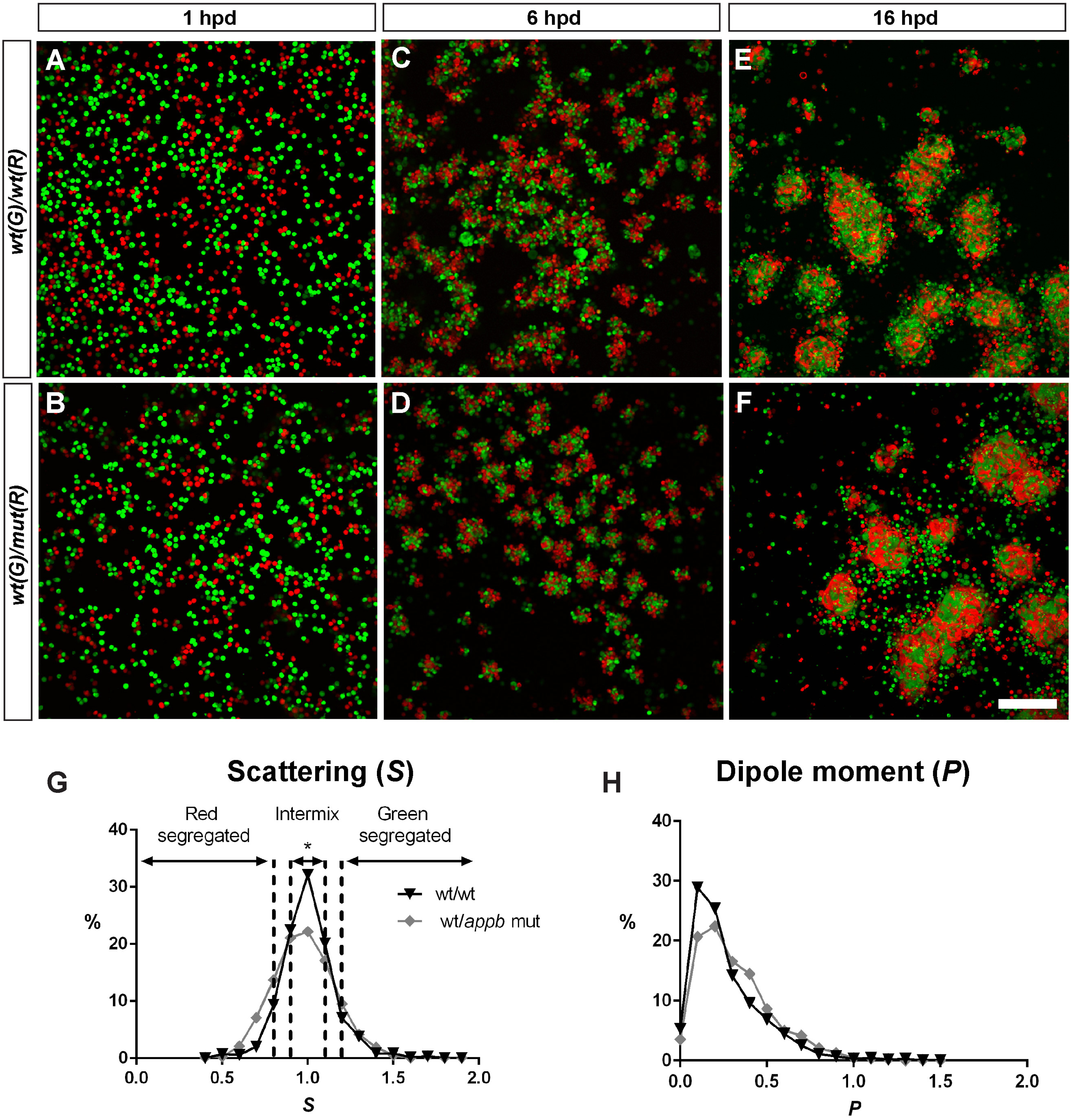
Appb regulates blastoderm cell adhesiveness. Embryos of wild-type or *appb* mutant background were injected with red (R) or green fluorescent dextran at the one-cell stage, dissociated and mixed 1:1 at sphere stage and co-cultured in fibronectin coated plates (A,B). Cluster formation started around 6 hours post dissociation (hpd) (C,D) and was allowed to proceed for 16 hpd before image analysis (E,F). Scale bar=200μm. Frequency distributions of scattering ‘S’ (G) and dipole moment ‘P’ (H) of red and green cells in clusters of wt/wt (black line) and *wt*/*appb* mut (grey line). Dashed lines mark interval for scattering. * p<0.05.

When applying these parameters to our data, we found a significantly lower level of intermixed (0.9>S< 1.1) clusters in wt^(G)^/mut^(R)^ (43%) compared to wt^(G)^/wt ^(R)^ clusters (56%, p<0.05), as shown in the frequency diagram (Fig. 6G). In addition, the shape of the S-curve has a tendency of being broader, suggesting a higher percentage of low S-values in clusters with mutant cells. This, in combination with the tendency of P being shifted towards 1 in wt^(G)^/mut^(R)^ clusters, shows increased segregation in clusters with wt and *appb* mutant cells and that these even may separate from each other, supporting a role of Appb in cell adhesion.

### No change in expression of tight- or adhesion junction markers ZO-1, E-cadherin and β-catenin

The above results led us to examine the arrangement of the cell-cell junctions. We stained embryos at 5.7 hpf for ZO-1, a protein expressed at tight junctions, and E-cadherin, a protein present in adhesion junctions. However, we were not able to detect any differences in neither the level or distribution pattern of either protein between wild-type (Fig. 7A,C) and *appb* mutants (Fig. 7B,D). As E-cadherin establish connections with the cytoskeleton through binding β-catenin (Aberle et al., 1994; Ozawa and Kemler, 1992), we also analyzed the expression of β-catenin in embryos at 5.7 hpf and found no obvious change between wild-type and mutant (Fig. 7E,F). Together, these results suggest that the general structures of both tight and adherens junctions are intact and that E-cadherin, ZO-1 and β-catenin do not seem to be affected by the mutation in *appb*.

**Fig 7.**
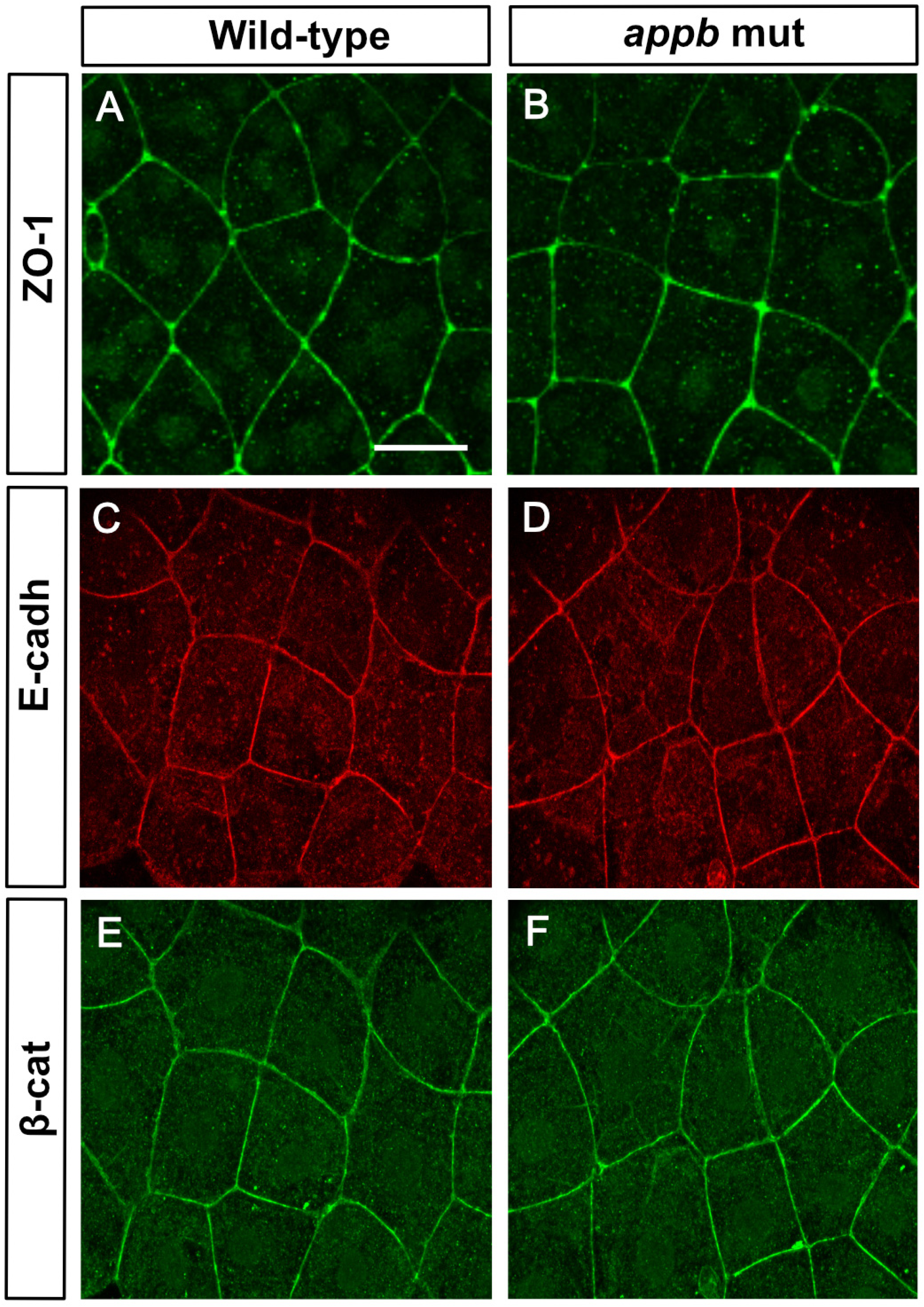
Normal expression of ZO-1, E-cadherin and β-catenin at 5.7hpf in *appb* mutants. Wild-type and mutant embryos at 5.7 hpf stained with ZO-1 (A,B), E-cadherin (C,D) and β-catenin (E,F). Wild-type (*n*=11) and *appb* mutant (*n*=12). Scale bar = 25μm

### Upregulated transcription of other App-family genes

The apparently normal phenotype of *appb* mutants at later stages indicate activation of compensatory processes. To investigate if the expression of other *app* family members was upregulated, we performed whole mount *in situ* hybridization and qPCR on wild-type and *appb* mutants for mRNA expression of *appa, appb, aplp1* and *aplp2.*

At 24 hpf, *appa* expression is found in tissues of mesodermal origin, lens, otic vesicles and somites (Musa et al., 2001). We observed a general increase in the expression of *appa* mRNA in *appb* mutants compared with wild-type embryos (Fig. 8A, upper panel). When we analyzed the expression closely at the head and trunk, we found that *appa* mRNA was highly increased in the brain (Fig. 8A, middle panel) and somites (Fig. 8A, lower panel).

**Fig 8.**
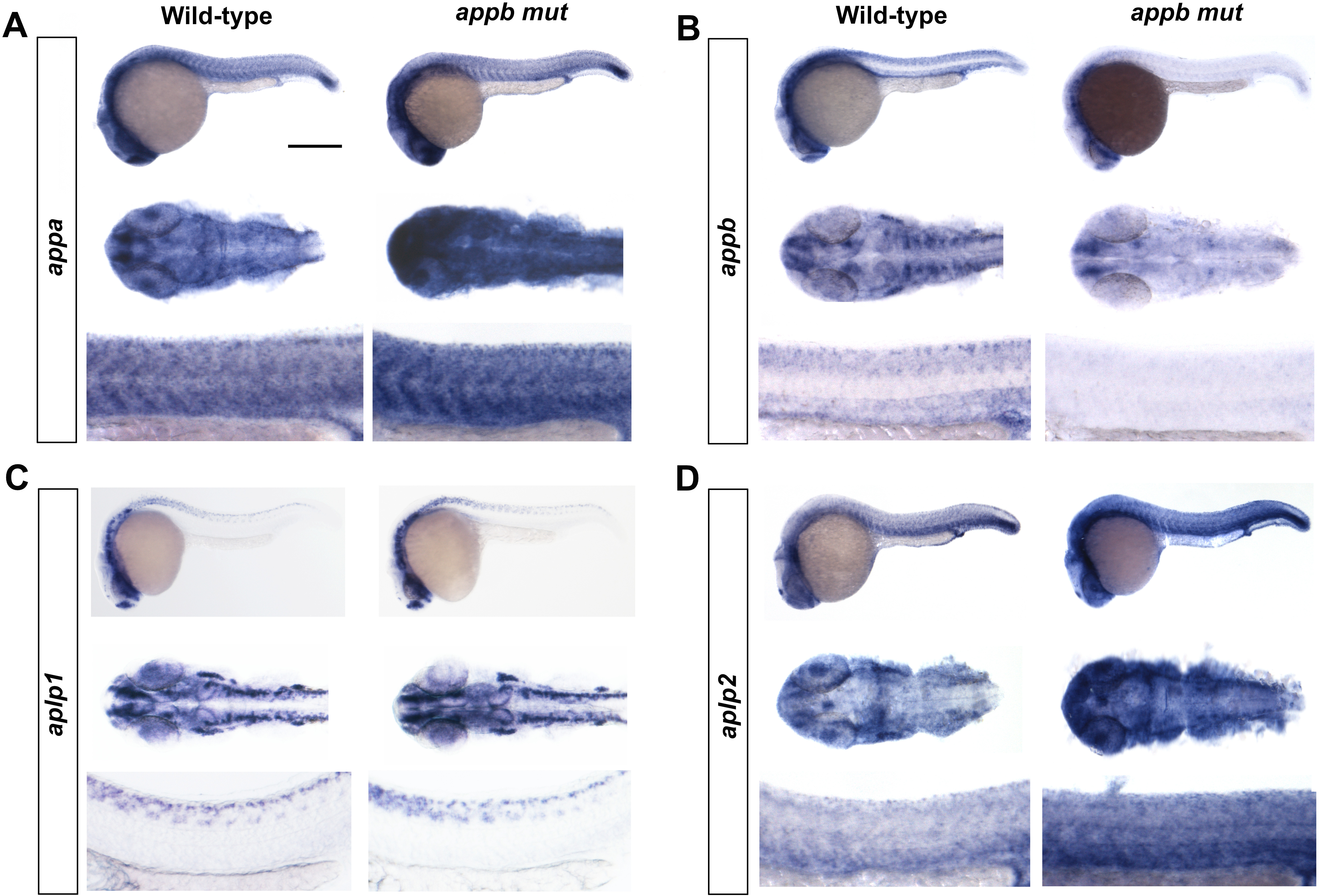
Expression of app family in *appb* mutants. Whole mount *in situ* hybridization of *appa* (A), *appb* (B), *aplp1* (C) and *aplp2* (D) at 24 hpf. Upper panel: whole embryos; middle panel: head (dorsal view) and lower panel: trunk in each set of genes (anterior to left). Scale bar = 500 μm (upper panel), 100 μm (middle panel) and 50 μm (lower panel).

The expression of *appb* transcript is localized to axial structures and is strongly expressed in neural tissues including telencephalon, mesencephalon hindbrain, trigeminal ganglia and spinal cord, as well as in pronephric duct and dorsal aorta (Abramsson et al., 2013; Musa et al., 2001). Prominent expression of *appb* mRNA is found in hindbrain rhombomeres 1-7, in clusters of cells organized in a ladder-like pattern (Banote et al., 2016). We found that the expression of *appb* mRNA was drastically reduced in *appb* mutants compared with their wild-type siblings (Fig. 8B, upper, middle and lower panel). In particular, nearly no expression was observed in the hindbrain and spinal cord region (Fig. 8B, middle and lower panel), confirming that the mutation results in increased degradation of *appb* mRNA.

The *aplp1* and *aplp2* genes are not well studied in zebrafish. Whole mount *in situ* hybridization revealed that *aplp1* was expressed in sensory neurons of the developing brain and spinal cord neurons (Fig. 8C). Strong expression was observed in the telencephalon, mesencephalon, adjacent hindbrain boundaries and trigeminal ganglia. No change in *aplp1* mRNA expression was detected in *appb* mutants (Fig. 8C, upper, middle and lower panel). In contrast, *aplp2*, with an mRNA expression pattern similar to *appa*, was also increased in *appb* mutants as compared to the wild-type embryos (Fig. 8D, upper, middle and lower panel).

Quantification of the expression of *app* family members in whole embryos by qPCR correlated well with the *in situ* hybridization results. In fact, the expression of *appb* was decreased (Fig. S2, B), while both *appa* and *aplp2* showed a tendency towards increased mRNA expression in *appb* mutants compared with the wild-type embryos (Fig. S2, A, D). The *aplp1* expression remained unchanged in the mutants (Fig. S2 C).

This led us to address protein expression of Appa and Aplp2 in mutants using the Y188 APP antibody that recognizes Appa, Appb and Aplp2. Western blot on whole embryos at 24 hpf showed increased Appa and/or Aplp2 protein levels in *appb* mutants compared with wild-type embryos (Fig. S2 E,F).

In summary, these results show that the *appb* mutation result in loss of Appb, which is accompanied by increased expression of Appa and/or Aplp2, that potentially could compensate for the deficits seen in mutants at early developmental stages.

### Normal locomotion at larval stage

To investigate whether the absence of Appb protein leads to behavioral defects, we monitored locomotor activity in the zebrafish larvae. To that end, we bred F3 heterozygous parents to generate F4 offspring. Comparison of the locomotor activity of the wild-type and *appb* mutants at 6 dpf revealed no significant difference in the distance traveled by *appb* mutants and wild-type larvae recorded over a 60-min time frame (Fig. 9A). In addition, the assessment of distance traveled at different velocity [high (>6.1 mm/sec), medium (2.1-6.1 mm/sec) and slow (<2.1 mm/sec)] did not show significant difference between these two groups (Fig. 9B-D).

**Fig 9.**
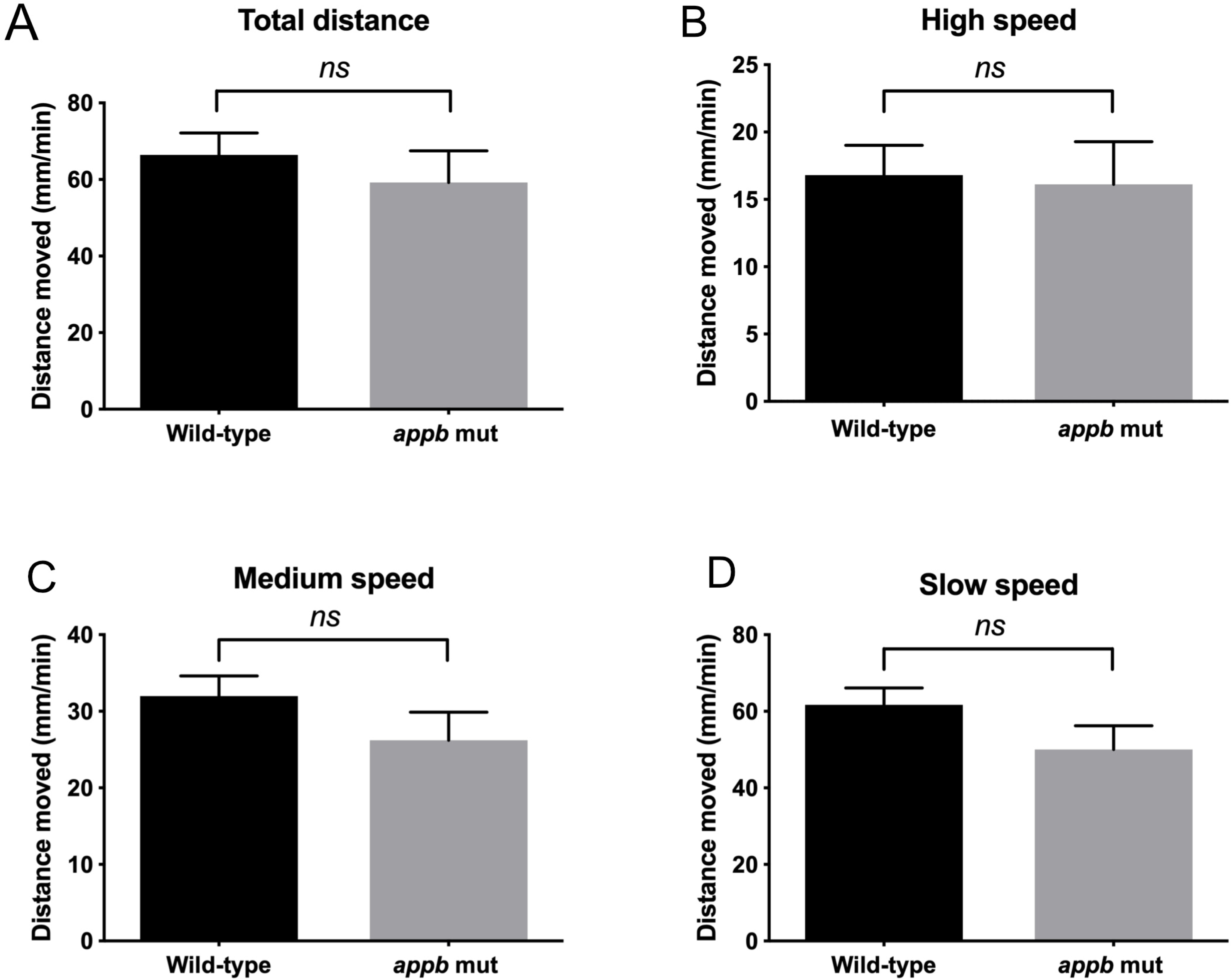
Normal locomotor activity in *appb* mutants at 6 dpf. Swimming activity over 60 minutes displayed as total distance traveled per minute (A), or as distance traveled at high (B, <6.1mm/s), medium (C, 2.1-6.1 mm/s) and low (D, >2.1mm/s) velocity. Wild-type (n=30) and *appb* mutant (n=23). Data were analyzed using Student’s *t*-test.

## Discussion

In the present study, we describe zebrafish with a homozygous mutation that introduces a premature stop in exon 2 of the *APP* homologue, *appb*. Similar to *App* mutant mice (Müller et al., 1994; Zheng et al., 1995), zebrafish lacking *appb* were initially smaller but morphologically normal, healthy and fertile as adults. Due to the external development of zebrafish embryos, we were able to identify a phenotype previously not described, which indicates a role of *appb* in cell adhesion during early development. The physiological function of APP family members has been hard to discern, likely due to the apparent redundancy between the APP and APLP2. As zebrafish have two *APP* homologues, *appa* and *appb*, in addition to the *aplp* genes, we expected *appb* mutants to have a similar or even milder phenotype than that described in mice. Indeed, this seems true after gastrulation. However, during early development, disrupted blastula integrity, together with general changes in EVL morphology and delayed gastrulation, clearly indicates the importance of Appb in these processes.

During the initial phase of gastrulation, the EVL is suggested to control thinning and extension of the underlying blastoderm cells, a process known as doming, through adjustments of surface tension (Morita et al., 2017). This process, as well as later stages of gastrulation, depends on tightly controlled cell-cell adhesion both within the EVL and in contact with the underlying blastomeres. Defects in EVL integrity are typically seen in zebrafish with genetic mutations affecting adherens junction, tight junction and cytoskeletal proteins (Eno et al., 2016; Lepage et al., 2014; Miles et al., 2017; Schepis and Nelson, 2012; Slanchev et al., 2009). Here we find that Appb is important for the cellular rearrangements during gastrulation. Although only a small fraction of *appb* mutant embryos displayed visible cell protrusions, their general sensitivity to additional destabilization of cadherin-mediated cell adhesion, indicate that cell-cell interactions are indeed weaker in mutants. Accumulating evidence shows that E-cadherin-mediated adhesion is crucial for proper gastrulation (Arboleda-Estudillo et al., 2010; Babb and Marrs, 2004; Kane et al., 2005; Shimizu et al., 2005; Song et al., 2013b). Interestingly, our *appb* mutants shared several phenotypes with E-cadherin hypomorphs (*hab*^*rk3*^), including uneven blastula surface, detaching cells and delayed epiboly (Shimizu et al., 2005). In addition, Appb knockdown experiments report convergence/extension and sensory ganglia defects similar to *hab*^*rk3*^ (Abramsson et al., 2013; Joshi et al., 2009). However, the lack of apparent changes in adhesion molecules (E-cadherin and β-catenin) or tight junction proteins indicates that Appb might control adhesion directly or through other pathways. Furthermore, the cell reaggregation experiment revealed that Appb controls cell-cell adhesion. These findings challenge previous results that did not reveal any difference in cell-cell adhesion between mouse embryonic fibroblasts from *App*-/- and wild-type mice (Soba et al., 2005). This discrepancy might further highlight the advantage of our model, since the *appb* mutant phenotype observed here likely appears before cells adapt to the loss of Appb. Thus, our data support that APP may act as a cell adhesion molecule. This is to our knowledge the first time an APP homologue has been shown to be crucial for cell-cell adhesion during early development.

Cell protrusions and EVL arrangements into rosette-like structures were characteristic for *appb* mutants. The EVL is subjected to stress during epiboly and to reduce tension anisotropy, the EVL normally undergoes oriented cell divisions (Campinho et al., 2013). However, although the larger size of EVL cells may reflect an increased epithelial thinning, our data did not show any significant difference in EVL cell number or proliferation. Thus, the lack of cell loss in the EVL made us hypothesize that the rosette-like structures observed in *appb* mutants might form as constriction points around EVL gaps arising from protruding cells.

An alternative explanation would be that the weak cell-cell adhesion between EVL cells force them to rearrange to release tissue tension. Rosette-like formations are transient structures formed by epithelial cells in various tissues including the lateral line primordium, pancreas, neuronal stem cells and epithelium and retina in *Drosophila* (reviewed by (Harding et al., 2014)). Such rosettes are often intermediate states during cell rearrangements and can be of two types, apical or planar polarized constrictions. In the EVL, cells are conformed in a planar polarized constriction, similar to that present during tissue elongation of the *Drosophila* epithelium (Blankenship et al., 2006). However, further studies are needed to elucidate the cause behind rosette formation.

The fact that most embryos with blastula defects managed gastrulation without major defects was surprising. In mice, the mild phenotype of APP-/-has been explained by redundancy with APLP2. Our results show that the loss of Appb correlates with upregulated expression of both Appa and Aplp2. It is thus likely that Appa and Aplp2 may compensate for the loss of Appb to support normal development after gastrulation. In mice, redundancy effects have been suggested since combined mutations in APP family member genes result in increased severity. However, reports on changed expression of APLPs at RNA or protein level are conflicting with some reporting no change in the APP-/-mice (Heber et al., 2000; Müller et al., 1994; von Koch et al., 1997; Zheng et al., 1995), while others show upregulation of both APLP1 and APLP2 in APP-/-mouse brain at 8 months of age but not earlier (Soba et al., 2005).

Recently, studies showed that mutated mRNA, especially if degraded as here, may bind and activate other genes (El-Brolosy et al., 2019; El-Brolosy and Stainier, 2017; Rossi et al., 2015). Interestingly, the degraded mRNA may preferably bind and upregulate closely related genes, which could explain many of the phenotypic discrepancies observed between knockdowns (using morpholino-injected embryos) and knockouts (genetic mutation) targeting the same gene. This mechanism was found activated not only in zebrafish, but also in fly and mouse (El-Brolosy et al., 2019). It is thus likely that degradation of mutated *appb* mRNA upregulates *appa* and *aplp2*, which consequently may diminish the more severe phenotypes observed in *appb* morphant compared with *appb* knockout fish (Abramsson et al., 2013; Banote et al., 2016; Joshi et al., 2009).

Modulation of APP expression results in cellular changes that may be unveiled as functional changes in locomotor behavior. In mice, loss of App leads to reduced locomotor activity (Guo et al., 2012; Zheng et al., 1995), whereas APP overexpression results in hyperactive behavior in adult mice (Rodgers et al., 2012). In contrast, *appb* mutant zebrafish embryos showed no significant change in behavior at 6 dpf. This discrepancy could be explained by the fact that rodent behavioral studies are performed on adults while the zebrafish behavioral responses in this study were monitored during early development. However, we cannot exclude that compensatory mechanisms of Appa and Aplp2 upregulation may contribute as well.

In summary, we conclude that Appb controls cell-cell adhesion at least during early development. This is, to our knowledge, the first time an APP homologue has been implicated in cell adhesion processes *in vivo*. Our data also shows that while loss of Appb is partly lethal, other related App-family members may create a back-up system to compensate for any loss of function. How these results translate to cell interactions and homeostasis in adult zebrafish and to APP function in mammals remain a subject of study. However, as mechanisms regulating cell adhesion during development often are involved in maintaining tissue organization and homeostasis in adults (reviewed in (Batlle and Wilkinson, 2012)), it is likely that the physiological function of Appb in adhesion is maintained throughout life.

## Acknowledgements

We thank Elisa Alexandersson for fish maintenance, Joachim Sturve for providing behavioral equipment and Alexandra Schauer for sharing protocols on cell dissociation and reaggreagation. We also acknowledge the Centre for Cellular Imaging core facility at the University of Gothenburg, the National Microscopy Infrastructure (NMI) (VR-RFI 2016-00968) for support with image acquisition and image analysis and the Genome Engineering Zebrafish facility (SciLifeLab, Uppsala, Sweden) for generation/housing of mutants.

## Author contributions

Conceptualization: A.A.; Methodology: A.A., R.K.B., J.C, T.M.S., R.C.; Software: R.C.; Formal analysis: A.A., R.K.B, J.C.; Investigation: R.K., J.C., T.M.S., A.A., G.V.K, Resources: J.L., G.V.K, S.M.B.; Writing - original draft: R.K.B., A.A.; Writing - review & editing: R.K.B., A.A, J.C., H.Z; Visualization: R.K.B., A.A., J.C.; Supervision: A.A, H.Z, S.M.B.; Funding acquisition: H.Z, S.M.B.

## Data availability

Description on image analysis of cell aggregation analysis will be available at https://github.com/CamachoDejay/.

## Supplementary data

### Supplementary information

**Supplementary data 1: Aggregation image analysis.**

**Fig S1.**
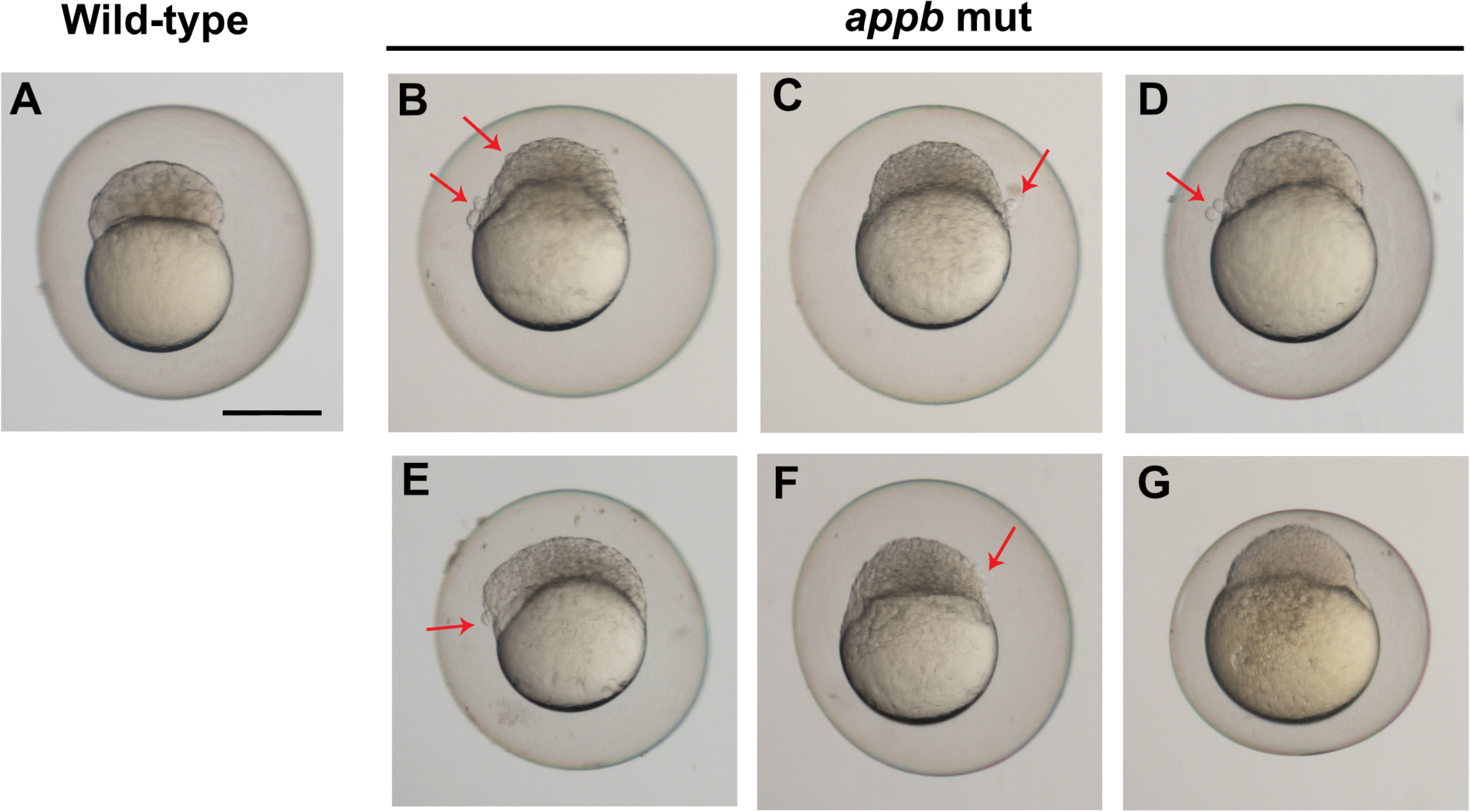
Blastula phenotypes in *appb* mutants. Examples of *appb* wildtype (A) and *appb* mutant (B-G) blastulas displaying clusters of cells budding from the EVL (arrow) and perched blastomeres resulting in an uneven blastula formation. Scale bar, 500μm.

**Fig S2.**
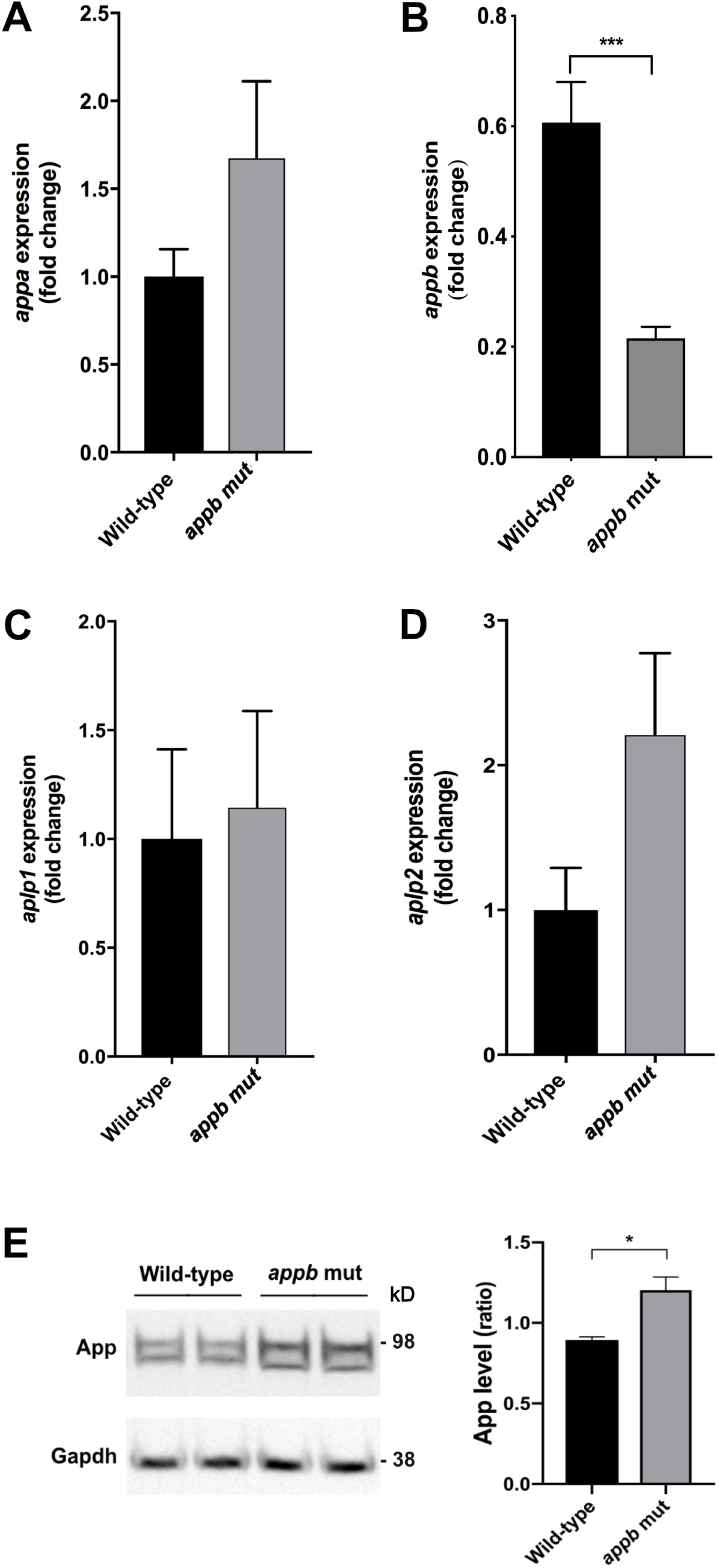
qPCR and WB analysis in *appb* mutants. Whole embryo qPCR analysis of *appa* (A’), *appb* (B’), *aplp1* (C’) and *aplp2* (D’) at 24 hpf. Values are reported as mean±SEM. (E) Western blot analysis and quantification of zebrafish App expression at 24 hpf, *n*=4. \**p*<0.05, \*\*\**p*<0.001.

## Supplementary data1

### Details on Image Analysis - aggregation and cell cohesiveness assay

This document will summarize:

1. How the algorithm for aggregate segmentation works.
2. How the calculation of the parameters S and P are done from the ROI images of a segmented aggregate.1.

1. **Aggregate segmentation pipeline – ROI generation**
  a. Load “.nd2” file. Image data is treated as a matrix of dimensions x,y,z, where x is the number of row pixels, y is the number of column pixels and z is 2 - the number of color channels (red and green).
  b. Calculate maximum intensity projection over the z dimension, effectively merging both color channels.
  c. Image contrast is enhanced via the Normalize Local Contrast method - FIJI
  d. Further improvement is done via a Gaussian blur – sigma 1
  e. Image thresholding via Li’s Minimum Cross Entropy method
  f. Objects smaller than 1000 pixels are removed from the segmented image
2. **Calculations of P and S for each ROI** Calculations are based on the article: Schötz, E., Burdine, R. D., Jülicher, F., Steinberg, M. S., Heisenberg, C., & Foty, R. A. (2008). Quantitative differences in tissue surface tension influence zebrafish germ layer positioning. HFSP Journal, 2(1), 42–56. https://doi.org/10.2976/1.2834817

#### P - closely related to the electrical dipole moment

Each image channel, red and green, are loaded as matrices, where the matrix value indicates intensity and the row and column index represent the pixel position. The only difference between the calculations for the red and green channel is that the pixel values of the red channel are considered negative, again in analogy to electrical charges.

For each channel, with intensity matrix *I*(*r, c*), where r and c are row and column indices, respectively:

1. Image is normalized, so the total sum on the pixel intensities is 1.

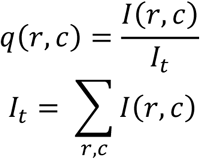
2. In an analogy to the dipole moments, the moment of each pixel 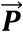 is calculated according to:

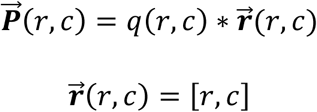
3. Then the total moment calculated by summing the moment of all pixels:

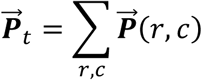

To allow for the comparison of aggregates of different size, 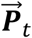 are normalized to the radius of the aggregate. This is done by:

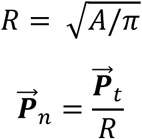

Where, *A* is the area of the aggregate in units of pixels.

Finally, the parameter P is calculated by summing the red and green moments and calculating the norm of the resulting vector:

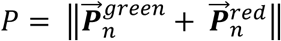

#### S - closely related to the moment of inertia and ratio of scattering amplitudes

Each image channel, red and green, are loaded as matrices, where the matrix value indicates intensity and the row, column indices represent the pixel position.

For each channel, with intensity matrix *I*(*r, c*), where r and c are row and column indices, respectively:

1. Image is normalized, so the total sum on the pixel intensities is 1.

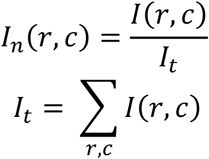
2. The center of mass of the image is calculated:

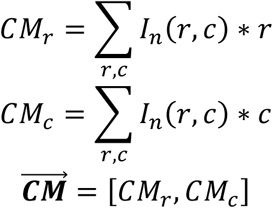
3. Calculate the displacement of each pixel from the center of mass:

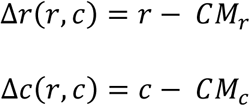
4. Calculate the tensor of inertia according to:

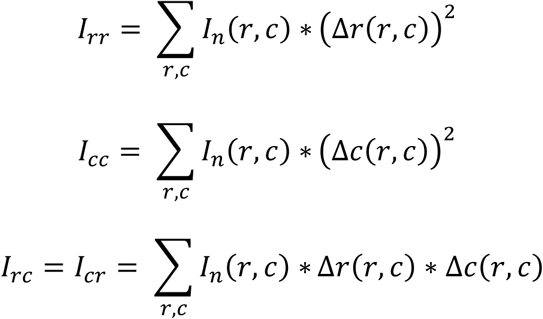
5. Calculate the principal moments of inertia:

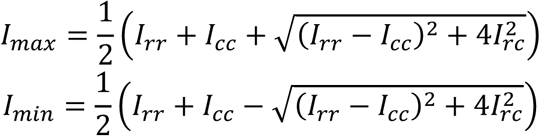
6. Calculate the scattering amplitude:

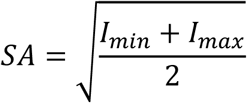

Finally, the parameter S is calculated as the ratio of the scattering amplitudes:

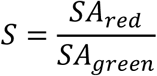

S is <1 if the red cells are less scattered, and >1 if the red cell are more scattered around the center of mass than the green cells. S=1 if cells are scattered equally.

